# Triadic Dynamics of Gastric Bacterial Microbiome, Phageome, and Host Genotype, with Implications for Disease Associations

**DOI:** 10.1101/2024.06.16.598898

**Authors:** Aleksander Szymczak, Katarzyna Gembara, Joanna Majewska, Paulina Miernikiewicz, Stanisław Ferenc, Jan Gnus, Marlena Kłak, Natalia Jędruchniewicz, Wojciech Witkiewicz, Krystyna Dąbrowska

## Abstract

The microbiota plays a key role for human health, and microbiome composition has been linked to specific health disorders many times. Different fractions of the microbiome enter dynamic interactions, particularly bacteria and bacteriophages (phages) – viruses preying on bacteria. Since the microbiome exists inside the human body, the body strongly affects the microbes, although not in a uniform way, but shaped by the extreme diversity of human genetic variants. These triadic dynamics of the bacterial microbiome, phageome, and human host genotype remain poorly understood; hence our goal in this study was to comprehend them as a holistic and interdependent system.

Gastric biopsies were the source of the stomach microbiome (bacteria and phages). Genotyping of patients was conducted on blood samples. They were analyzed by next generation sequencing followed by multiway statistical comparisons of identified bacterial and phage taxa, human genetic variants, and medical data from the patients, including gastric disorder diagnostics.

Commonly expected associations between presence of bacteriophages and their specific hosts were not found to be a universal principle. With comparable SNP correlations, the strongest associations between *Staphylococcus* and some staphylococcal phages or *Enterobacteria* and some enterobacteria phages were discovered. However, many more phage groups were not found to be clearly associated with their bacterial hosts, though often associated with SNPs, particularly those linked to immunological functions and body responses to bacteria and viruses. Thus, in addition to the expected effect on phage communities by shaping the communities of their bacterial hosts, the human body seems to affect phages directly, selecting for phages that survive its specific selective pressure, which can be defined by detection of particular genetic variants.

## Introduction

Each human body is a unique host for 10-100 trillion microorganisms. Shotgun sequencing revealed almost 3.3 million non-redundant genes in the human gut alone. Despite coding regions that could be detected in other parts of the body, this is over 150 times more than the number of genes in the entire human genome (approximately 22 000) (Abdellah et al., 2004; Qin et al., 2010). Observations demonstrating the key role of the microbiota for human health have induced wide interest in the composition of the human microbiome (Fan & Pedersen, 2020; Gill et al., 2006; Peterson et al., 2009; VanEvery et al., 2022). The bacterial part of the microbiome can be analyzed by 16S rRNA (a small subunit of bacterial ribosomes) gene sequencing, since bacteria possess a characteristic variant of this gene due to its hypervariable regions. This has allowed for extensive studies of the bacterial fraction of the microbiome, with increasing evidence for its key role in health and disease (Prodan et al., 2020). In turn, the viral fraction of the microbiome, particularly bacteriophages, which in fact outnumber bacteria and other microbes, have no simple genetic signature that might help in fast and easy identification of their complex taxonomy in body niches. Shotgun sequencing remains the major tool in studies of viral communities (Castelino et al., 2017; Ranjan et al., 2016). Of note, both methods also detect microorganisms impossible to culture and detect using traditional laboratory techniques, and they can give an insight into true microbiome composition.

The human body includes various environments (niches) for microbes, where conditions are different; one of the major microbial communities in the human body resides in the gastrointestinal tract. Depending on the part of this system, we can identify different communities of bacteria living there. In particular the intestine has been demonstrated as the major “body bioreactor” with a considerable microbial mass. The stomach on the other hand was long considered sterile because of unfavorable physico-chemical conditions in this organ. Gastric juice makes it extremely hostile to any life forms, since it contains hydrochloric acid and pepsin (endopeptidase), with pH reaching 1.7 (Dressman et al., 1990). However, some microbes - including viruses - can create a core community that inhabits that organ. Rare studies have revealed the presence of *Firmicutes*, *Bacteroidetes*, and *Actinobacteria* in gastric mucosal samples. One of the most recognizable species is *Helicobacter pylori* – an important pathogen that strongly increases the risk of stomach diseases (Nardone & Compare, 2015b; Z. Wang et al., 2020; Y. Yang et al., 2021). Lactic acid bacilli that have also been identified as a probiotic part of the gastric microbiome have also developed the ability to survive in the unfavorable environment of gastric pH (Deckers-Hebestreit & Altendorf, 1996). *Escherichia coli* has evolved a unique ability to avoid negative effects of low pH in the stomach by rearrangements of lipids in the outer membrane (Foster, 2004; Lund et al., 2014; Y. Xu et al., 2020). A part of the microbiota found in the stomach was identified as transient and related to a diet, for instance pathogenic *Salmonella*, or to the oral cavity, e.g. *Streptococcus* They often demonstrate some adaptations to lower pH, but commonly bacteria take advantage of the fact that the pH of gastric juice varies. After a meal, the pH temporarily rises, and this also provides an opportunity for the bacteria to have a limited hold on the stomach mucosa (Nardone & Compare, 2015b; I. Yang et al., 2013)

The stomach microbiome also includes a numerous group of viruses, mostly bacterial viruses. Their adaptations to the stomach environment are poorly known. Specific factors coded by phage genomes as adaptations to the stomach environment have been demonstrated so far only in T4-like phages, where the structural protein gp24 mediates phage resistance to low pH and digestive enzymes (Majewska et al., 2019). Within most investigated phages, many have been demonstrated inactive when pH in the environment drops below 4. Some exceptions, such as M13 or T1 phages specific to *Escherichia coli*, can survive pH as low as 2 and 3, respectively. In addition to virion forms, bacteriophages also have the ability to survive harsh environmental conditions inside bacterial cells as an incorporated part of bacterial genomes (prophages), or in a plasmid-like form, if able to enter life cycles other than the lytic one (Jończyk et al., 2011; Nobrega et al., 2016).

Bacterial and bacteriophage communities are interdependent, since bacteriophages that prey on bacteria control bacterial populations, and at the same time phages need bacterial hosts to propagate. Further, this interdependent community exists inside the human or animal body (mammalian host), which may demonstrate individual features, particularly determined by individual genetic characteristics. The complex body environment in which microbiota exist is also shaped by mammalian genetic determinants. Human genomes may include 4.1 to 5 million different variants, more than 99% of which are single nucleotide polymorphisms (SNPs) and indels. Next generation sequencing (NGS) techniques have revealed associations between some human genome variants and profiles of the human microbiome but only in other (than the stomach) body niches, (Igartua et al., 2017a, S. Lee et al., c2011, Awany et al., 2019b, Dabrowska & Witkiewicz, 2016). On the other hand, observations that negatively verified hypotheses on microbiota-human genome association have also been reported (J. Wang et al., 2016).

Data on the stomach microbiota are still very limited; in particular, the stomach phageome has not been described so far. In this study we sought to comprehend the bacterial microbiome, phageome, and host genetics as a holistic and interdependent system. We aimed to identify three-way associations among the bacterial fraction of the stomach microbiome, stomach bacteriophages, and the genetic background of the mammalian host. Additionally, we sought to ascertain whether certain bacterial and viral elements of the stomach microbiome are linked to gastric diseases.

## Materials and methods

### Bioethical statement

Samples were collected in accordance with the principles of good clinical practice and the Declaration of Helsinki. The research was approved by the local Bioethical Committee of the Regional Specialist Hospital in Wroclaw (approval no. KB/nr 8/rok 2017) During the individual interview, all information about the study was provided and written consent was obtained from each participant.

### Sample collection from human participants of the study

Gastric biopsies used in the experiments were taken by qualified physicians working in a Regional Specialist Hospital in Wroclaw, Endoscopy Department. Samples included biopsies collected during gastroscopy examination by the physicians. Prior to the examinations, patients consented to the use of biological samples collected from them, and were interviewed on chronic diseases, medications they were taking, and on antibiotics use (if antibiotics had been used within the last 6 months, the patient was not included in the study). The biopsy was taken from the pyloric part of the stomach. That specific location was chosen according to relevant literature identifying it as the part of human stomach most inhabited by microorganisms; this area is also recommended for diagnostics of *Helicobacter pylori* infection (Bayerdorffer et al., 1989). In addition, a blood sample was collected from a patient for further SNP analysis. Of note, in some patients blood samples were not available, or parallel isolation of the bacterial and bacteriophage fraction of the microbiome from biopsies was not possible due to technical issues. Relevant information on the number of available samples is given for each type of analysis (see below: N for each library type).

Each patient was diagnosed according to the International Classification of Diseases (ICD) (Drösler et al., 2021); a full list of patients’ ICD codes is presented in Table S1. Patients’ demographic and medical data were derived from the interview and from the medical database Asseco Medical Management Solutions (AMMS). The data included: age, gender, chronic diseases affecting the patient, medications taken permanently, recent (6 months) history of antibiotic treatments, and drugs from the group of proton pump inhibitors. Numpy was used for array analysis in numerical data (Harris et al., 2020). Scikit-bio – a python package with algorithms, and resources for bioinformatics – was used for index calculations.

### DNA isolation for NGS

Each biopsy collected from a patient immediately after collection was placed in PBS (2 ml) and forwarded for separate viral and bacterial DNA isolation. Specimens in PBS were floated on a 3D shaker for 3 h at 4C and centrifuged at 12 000g for 10 minutes. The supernatant was used for further bacteriophage separation and the pellet was subjected to bacterial DNA isolation with the Micro Beat Bead Gravity AX kit (A&A Biotechnology). Bacteriophage separation was performed with cesium chloride (CsCl) ultracentrifugation as follows: the supernatant was filtered through 0.22 μm pore membrane and then loaded on CsCl gradient. The CsCl density gradient ranged from the highest density, 1.7 g/ml, followed by 1.5 g/ml and 1.35 g/ml, to 1.15 g/ml. Centrifugation was performed overnight at 62 000 g at 4C in a swinging bucket rotor. Half a milliliter of content between 1.35 and 1.5 g/ml of CsCl solution was collected with a syringe for further phage DNA isolation. DNA isolation was performed with a Sherlock AX (A&A Biotechnology) kit. Samples were proceeded to Genomiphi V2 DNA Amplification (Cytiva). Blood was drawn with the intention of obtaining a clot. From it, human genomic DNA was isolated using Genomic Micro AX Blood Gravity Kit (A&A Biotechnology). DNA isolated from biopsy samples using the Micro Beat Bead Gravity AX Kit (A&A Biotechnology) was subjected to bacterial DNA sequencing. DNA samples isolated from human blood using Sherlock AX (A&A Biotechnology) were forwarded to Ampliseq – SNP sequencing. DNA obtained from biopsy samples and ultracentrifuged were subjected to Shotgun sequencing for phageome investigation.

### Library preparation for 16S rRNA sequencing (N=128)

Initially, bacterial DNA isolated in the previous step was quantified using Qubit 2.0 with the dsDNA High Sensitivity Assay Kit (Thermofisher). The minimum concentration of DNA that was necessary for further steps (no less than 2 ng/μl) was achieved in 128 samples; these samples were collected for further analysis. Amplification of 16S rRNA regions was performed with the Ion 16S Metagenomics Kit (Thermofisher). For sample pooling oligonucleotide barcodes were included in the library preparation; the Ion Xpress Barcode Adapters kit (Thermofisher) was used as recommended for enzymatic fragmentation along with the Ion Plus Fragment Library Kit.

### Library preparation for phageome sequencing (N=96)

Eluted viral DNA was quantified using Quantus Fluorometer with the QuantiFluor ds DNA system kit (Promega). Minimum concentration of DNA that was considered as sufficient was 1 ng/μl and samples with lower dsDNA concentration were excluded. In total, 96 samples met quality requirements to be processed for further sequencing. DNA was then amplified with the GenomiPhi V2 DNA Amplification Kit (Cytiva Life Sciences) and again quantified using the Quantus Fluorometer and QuantiFluor dsDNA Kit (Promega). Illumina DNA Prep (Illumina) was used to prepare sequencing libraries. All samples were barcoded using Nextera DNA CD indexes (Illumina). Final concentration of libraries was calculated using Quantus Fluorometer results.

### Library preparation for SNP sequencing (N=72)

Quantitation of human DNA isolated from blood cells was performed using Qubit 2.0 with the dsDNA High Sensitivity Assay Kit (Thermofisher A custom Ampliseq panel for the detection of SNPs was designed using Ion AmpliSeq Designer and ordered as Ampliseq Custom Panel, Thermofisher). It contained 212 SNP targets within 181 different amplicons (Supplementary Table 1). Multiplex PCR was designed using Ion Ampliseq Designer. For the amplification of dedicated regions Ion Ampliseq Library Kit Plus (Thermofisher) was used according to the manufacturer’s instructions. For later sample pooling oligonucleotide barcodes were introduced during the library preparation: the Ion Xpress Barcode Adapters kit (Thermofisher) was used. In total, an AmpliSeq library was prepared for 72 samples.

### IonTorrent sequencing

Quantified libraries were proceeded to emulsion PCR using Ion One Touch 2 according to the manufacturer’s instructions. Initialization was performed using the IonTorrent Hi-Q View Kit (Thermofisher) with Ion 318 Chip Kit v2 BC (Thermofisher Scientific). Sequencing was performed using the Ion Torrent Personal Genome Machine.

### Illumina sequencing

For final library preparation a NextSeq 500/550 Mid Output kit (Illumina) was used according to the manufacturer’s instructions. Sequencing was performed using a NextSeq 550 device with default settings and 2×150 reads.

### Files extraction

FASTQ files were downloaded from sequencing devices using dedicated software for Illumina and Ion Torrent devices, that is BaseSpace (Illumina) and TorrentExporter (Thermofisher Scientific) respectively. Files from IonTorrent PGM were exported as single end sequences; thus read pairing was not applicable. Data were transferred to an internal computer cluster in Hirszfeld Institute of Immunology and Experimental Therapy (HIIET).

### 16S rRNA data analysis

To create FASTQ files from 16S rRNA sequencing the FileExplorer plugin was used on the IonTorrent Server. Data were transferred to a local computer cluster in the HIIET. Firstly, FastQC – a quality control tool for High Throughput Sequence – was used to determine quality of performed runs. This tool was created for screening potential errors in datasets created using NGS technologies. To remove low quality sequences from the analysis, Trimmomatic V0.32 (Bolger et al., 2014) was used with a minimal length of 50 base pairs (bp). Sequence mapping was performed with Kraken2 (Wood et al., 2019). Kraken2 is a taxonomic classification tool which makes use of k-mer matching. The Kraken Microbial Database (September, 2018), which includes all 16S rRNA genes publicly available in the public repositories, was used.

### Viral data analysis

FASTQ files were obtained by demultiplexing. Data were transferred to a local computer cluster in the HIIET. FastQC was applied to detect possible low-quality files. Low-quality sequences were removed from the analysis by Trimmomatic V0.32 (minimal length of 50 bp). By mapping sequences to the Human Genome GRCh38 (GenBank Accession: GCA 000001405.15) and removing positive hits from further analysis, the considerable contribution of human DNA was eliminated from the analysis. This step was completed using bowtie2 (Langmead & Salzberg, 2012). Initially, unaligned sequences were processed using Kraken2 aligner, with the Kraken Microbial Database (September, 2018). However, due to dynamic changes in taxonomy annotations related to bacteriophages it turned out to be outdated. It was necessary to prepare a custom database and use another, more customizable aligner. Therefore, sequences unaligned to the reference human genome were subjected to BLAST analysis. The BLAST algorithm was used along with parameters that allowed 2 mismatches in an aligned sequence and a minimum length of 50 bp. BLAST was used locally on the computer at HIIET. The reference for this part of the analysis was built from all DNA virus sequences available in the public National Center for Biotechnology Information Genomes repository.

### Single nucleotide polymorphism AmpliSeq data analysis

FASTQ files were generated on the IonTorrent server and then exported using the FileExplorer plugin to a computer cluster in the Institute of Immunology and Experimental Therapy. Quality of the reads was assessed using FastQC and low-quality sequences were removed from the analysis by Trimmomatic V0.32 (minimal length of 50 bps). Burrows–Wheeler Aligner (BWA) (Li & Durbin, 2009) was used to align DNA sequences to a reference human genome: Human Genome GRCh38 (GenBank Accession: GCA_000001405.15) that was indexed with the “bwa index” command. Generated Sequence Alignment Map (SAM) files were converted to Binary Alignment Map (BAM) with SamTools version 1.14 (Li et al., 2009). BAM files were sorted, and duplicates were marked using Picard. It is a predefined set of command line (CLI) programs that were designed for processing files in formats such as SAM/BAM/VCF. BIM Collaboration Format (BCF) tools were designed to perform manipulations on Variant Call Format (VCF). This was the tool that was used to transform sorted BAM files to VCF as applicable for further analysis. This step is called variant calling, which refers to the identification of variants within sequenced data. VCF files were further processed with the SNPeff (Cingolani et al., 2012) tool, which identifies SNPs present in sequenced samples.

### Data curation in Pandas

Data were curated according to Mckinney (2010). Data derived from all 3 types of NGS data analysis were transformed into common formats for further statistical data analysis. For this purpose, each folder with respective data was merged into one comma-separated values (csv) file. 16SrRNA data before transformation were saved in kraken2 output files containing data on classified microorganisms, accuracy, and counts per taxonomy level. The Bash shell was used.

AmpliSeq sequencing yielded VCF files. Data such as a SNP’s reference, position, quality, and gene were derived from these data and merged into one csv file to make it readable for python libraries used for statistical data analysis. The Bash shell was used.

Data exported from BaseSpace Illumina sequencing and analyzed yielded numerous output files from the BLAST aligner. Those files were imported to the JupyterLab environment using Pandas (Python Data Analysis Library). The files contained information about aligned sequences and NCBI taxid numbers. To retrieve full taxonomy data, such as classification and nomenclature, for all the organisms annotated in public reference databases, the taxonomizr (0.9.3) package was used in the RStudio 4.1 environment. Its function is to allocate taxonomy to an NCBI accession number. This package made it possible to download data dumps (last updated on 26.02.2022) and to create a local repository for custom taxonomic assignment. R 4.1 was used.

### Statistical data analysis

Correlation between detected SNPs and the diversity index in 16S rRNA and bacteriophage elements of the microbiome was estimated using the Shannon-Wiener equation (Spellerberg & Fedor, 2003). Its value determines the probability that two random elements from a sample belong to different species. Taxonomy level of species was used to calculate the index value for bacteria. In order to assess possible associations between the genotypes of patients and the composition of their microbiome, the sequencing data for microbiome composition were converted into a binary format to standardize them with the SNP data format and to facilitate further analysis. A threshold of 10 reads (either positive or negative) was used in this conversion, with a taxon considered positive if at least ten reads were mapped. The Mann-Whitney test was employed to evaluate the significance of the correlation between the bacterial and bacteriophage components of the patients’ microbiomes and the SNPs in their genomes.

Principal component analysis (PCA) was performed using RStudio environment in version 3.1. For the purposes of analysis, the number of reads for each group was normalized to the total number of reads that were obtained in each sequenced sample. The following libraries were imported and applied: “dplyr”, “tibble”, “ggplot2”, “ggfortify”, “tibble”, “stringr”, “data.table”; PCA plots are projections of multidimensional data on a 2D plane. R 4.1 was used to create an individual script. Biplot figures were created using the “factoextra” library in R. Dimensions used in these figures were chosen based on their variance coverage from scree plots generated using the same library in R. PERMANOVA analysis was applied using the “adonis” library in R.

### Analysis of correlation between SNP distribution and occurrence of bacteria presence using Cramer’s V test

Cramer’s V test was used as a measure of association between two nominal variables, based on Pearson’s chi-squared statistic. This assumption is satisfied by the factors that we can specify in the binary system, where their occurrence is the number 1, and their absence is the number 0. For this purpose, contingency tables were created using Pandas DataFrames. Cramer’s V equation was transformed into the Python 3.6 function.

### Test of proportion

Correlation between bacterial and bacteriophage presence was calculated using the test for proportions based on the normal z-test. The function from the statsmodel library coded in Python 3.6 was used to compute the results. The formula used for calculation is presented in Equation 2. Quantitative data from microbiome composition sequencing were transformed into categorical where variables were defined as “present” and “absent” in relation to bacteriophages or bacteria.

## Results

### Correlated microbial groups identified in the stomach microbiome

Gastric biopsies were collected from patients of the Endoscopy Department, Regional Specialist Hospital in Wroclaw, and processed for isolation of bacterial and viral fraction isolation. The bacterial fraction was analyzed by sequencing of 16S rRNA coding regions, and the bacteriophage fraction was isolated in a cesium chloride gradient and analyzed by shotgun sequencing. Relative abundance of bacterial and bacteriophage groups in the stomach microbiome was identified and correlations between identified groups were assessed using Pearson correlation (Python 3.6). A full list of paired microbial groups with r>0.5 and p<0.05 is provided in Supplementary material (Table S2). Selected correlations are discussed below.

### 1. Bacteria–bacteria correlations

Significant correlations between bacterial genera were identified in the stomach microbiomes of the investigated patients (Table S2); particular focus was given to genera belonging to the ESKAPE group of pathogens and to *Escherichia* sp., due to their marked contribution to the problem of multi-drug resistant hospital infections. Interestingly, *Escherichia* correlated strongly with *Shigella* (0.97), *Yersinia* (0.89), and *Salmonella* (0.80) and moderately (0.65) with *Enterobacter*. Presence of *Klebsiella* in the stomach microbiome of patients correlated with existence of *Enterobacter* (0.95), *Yersinia* (0.86), *Providencia* (0.73), and *Morganella* (0.71) species. A moderate correlation was found for *Acinetobacter* and *Helicobacter* (0.59). Also, *Acinetobacter* was moderately associated with presence of *Enterococcus* (0.56). No significant bacteria–bacteria associations were observed for *Pseudomonas*. A moderate association between *Staphylococcus* and *Enterococcus* (0.55) was observed (Table 1 and Table S2).

**Table 1.** Correlations of ESKAPE pathogens and Escherichia sp. to other bacterial groups.

| *Enterococcus* | *Staphylococcus* | *Klebsiella* | *Acinetobacter* | *Pseudomonas* | *Enterobacter* | *Escherichia* |
| --- | --- | --- | --- | --- | --- | --- |
| *Acinetobacter* ($r > 0.5$)<br>*Staphylococcus* ($r > 0.5$) | *Enterococcus* ($r > 0.5$) | *Enterobacter* ($r > 0.9$)<br>*Yersinia* ($r > 0.8$)<br>*Providencia* ($r > 0.7$)<br>*Morganella* ($r > 0.7$) | *Enterococcus* ($r > 0.5$)<br>*Helicobacter* ($r > 0.5$) | NONE | *Klebsiella* ($r > 0.9$) | *Shigella* ($r > 0.9$)<br>*Yersinia* ($r > 0.8$)<br>*Salmonella* ($r > 0.8$)<br>*Enterobacter* ($r > 0.6$) |

### 2. Phage–phage correlations

We identified 49 genera of bacteriophages present in the patients’ stomach mucosa, in total. Among them, the most numerous were 4 genera: *Kayvirus*, *Punavirus*, *Lambdavirus*, *Inovirus*.

We observed clear correlations between some bacteriophage groups that prey on the same bacterial hosts. *Lambdavirus* bacteriophages specific for *Escherichia* were significantly associated with same-host viruses: *Tequatrovirus* (0.78) and *Punavirus* (0.78); independently of these, other phages predating on enterobacteria including *Escherichia* sp. were found correlated: *Pankowvirus, Lederbergvirus,* and *Oslovirus*. Bacteriophages specific to *Staphylococcus* were also related to each other: *Triavirus* was associated with *Phietavirus* (0.83), *Dubowvirus* (0.83), *Peeveelvirus* (0.83), and *Biseptimavirus* (0.83). Weaker correlation (0.55) was independently observed between *Kayvirus* and *Twortvirus* (Figure 1). Bacteriophages specific to *Streptococcus* – *Moineauvirus* and *Brussowvirus* – were also strongly correlated (0.80) (Table S2).

**Figure 1.**
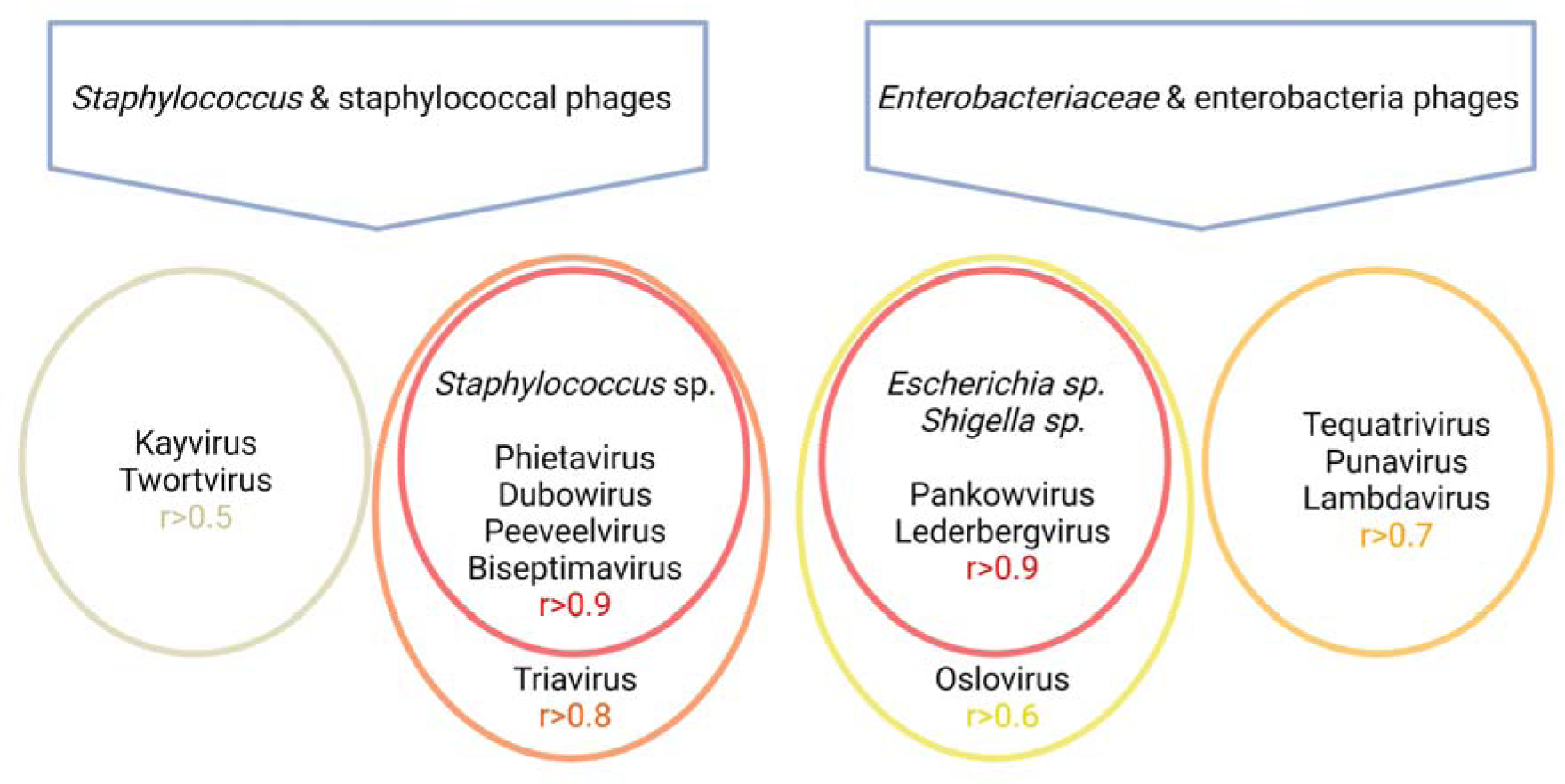
Major correlations between bacterial and bacteriophage groups found in human stomach microbiome.

### 3. Phage–bacteria correlations

Correlations between bacterial and phage components of the patient’s microbiome were tested. Interestingly, the same (correlated) bacteriophages specific to *Staphylococcus* such as *Triavirus, Phietavirus*, *Dubowvirus*, *Peeveelvirus,* and *Biseptimavirus*, were strongly correlated (r>0.9, p<0.001) with their potential hosts. Also, intercorrelated enterobacteria phages: *Pankowvirus*, *Lederbergvirus*, and *Olsovirus* were correlated to *Escherichia* sp. and *Shigella* sp. (Figure 1). These results demonstrate that some phage-host groups are correlated, while not all intercorrelated phages correlate at the same time to their host (like *Kayvirus* and *Twortvirus*, or *Tequatrovirus*, *Punavirus*, and *Lambdavirus*). Due to the diversity of phage life cycles, providing a definitive explanation for this phenomenon is challenging. However, we hypothesize that in phage groups that are intercorrelated with their hosts, the majority consists of temperate phages that function as prophages. On the other hand, among those phage groups that are intercorrelated independently of their bacterial hosts, there is a higher prevalence of obligatory lytic phage strains. (Figure 1). Surprisingly, we found no significant associations between *Pseudomonas*, *Klebsiella*, *Enterococcus*, *Enterobacter*, and *Acinetobacter* bacteriophages and their hosts (Table S2).

### Correlations of bacteriophages and their bacterial hosts with bacterial diversity (Shannon) in the stomach

Decreased diversity of microbiomes is a commonly known factor linked to dysbiosis and disfunctions of the gastrointestinal tract. Therefore, the analysis of a direct correlation between microbial groups was extended to correlations of microbial taxa to the overall microbiome diversity. We found 5 phage genera that were significantly associated with microbiome diversity measured by the Shannon index. Of those, the presence of the staphylococcal phage *Kayvirus* (p<0.05) and coliphages *Teseptimavirus* (p<0.05) and *Traversvirus* (p<0.05) was related to decreased diversity. However, other staphylococcal phages – *Dubowvirus* and *Phietavirus* – were significantly associated with increased diversity (p<0.05). *Teseptimavirus* is specific to *Enterobacteria*, *Escherichia*, *Salmonella* and *Yersinia* species. *Traversvirus* species include bacteriophages specific to *Escherichia* and Stx-converting bacteriophages; this suggests that in this case phages play a role as biomarkers; that is, they are overrepresented in patients affected with overgrowth of specific bacterial species. No other correlations between phage groups and the Shannon diversity index were found (Table S3).

We found 249 correlations among bacteria whose presence had a statistically significant effect on the diversity of the bacterial microbiome in the stomach. Surprisingly, *Helicobacter pylori* did not feature among these correlations (though it was commonly detected). This suggests that despite its importance for stomach health, in many patients *Helicobacter pylori* does not determine the overall composition of the stomach microbiome. Interestingly, the presence of *E. coli* and *Staphylococcus warneri* proved statistically significant for the diversity of the stomach microbiome according to the Shannon index (p<0.01); however, it was associated with increased diversity of the stomach microbiome in the investigated group of patients (Table S3), suggesting a positive implication for a healthy microbiome.

### Bacterial and phage groups associated with microbiome composition

PCA was used to identify bacterial and phage groups linked to differences in the overall composition of microbiomes. PCA was plotted using bacterial and phage groups identified within patients’ microbiomes. To evaluate crucial components for each cluster, differences between individual taxa were presented using the log2 fold change of normalized number of reads. The highest log2 fold change of the groups present in both clusters was observed among the phages *Ceduovirus* (specific to *Lactococcus sp.*) and *Sinsheimervirus* (specific to *E. coli*), and the bacteria *Desulfosporosinus*, *Acinetobacter*, *Christensenella*, *Achromobacter, Helicobacter*, *Oscillibacter, Pseudoxanthomonas*, and *Melissococcus*. PERMANOVA analysis revealed that the centroids of our clustered samples differ from each other significantly, with p<0.001. Interestingly, among other groups of bacteriophages that correlated with their hosts, there were no differences between the identified clusters. Results are presented in Figure 2.

**Figure 2.**
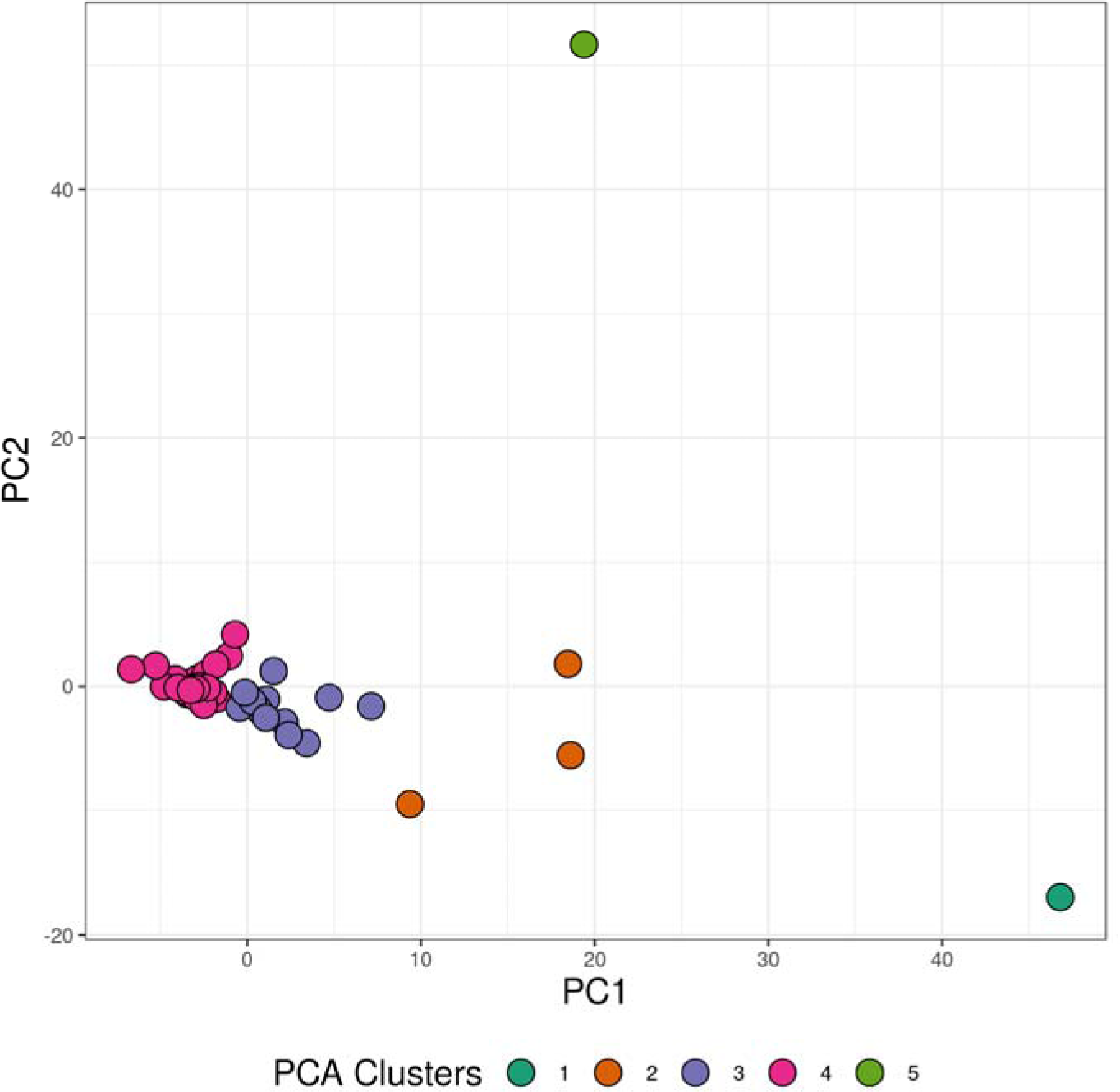
Interrelationships between bacterial and phage elements within the patients’ microbiomes identified by principal component analysis (PCA); significant differentiation between the centroids of Clusters 3 and 4 (p<0.001) was found.

### Bacterial and phage elements of the microbiome associated with gastric diseases

The diagnosis of each patient was made according to the International Classification of Diseases (Drosler et al., 2021); a complete list of patients’ ICD is given in Table S1. The one-proportion z-test was used to identify potential correlations between diagnosed gastric diseases and microbial groups detected in the patient’s stomach (patients with identified ICD-classified diseases were the ICD-positive group and patients without any diseases either diagnosed nor reported in the patient interview were the control group). The analysis revealed that 14 bacterial species correlated with ICD-classified diseases diagnosed in the investigated patients. Bacterial microbiome associations were found for diseases classified as: K29 *Gastritis and duodenitis*, R.10.4 *Other and unspecified abdominal pain*, K21 *Gastro-oesophageal reflux disease* (correlations defined by CI 0.95 and p<0.05) (Table 1). The analysis further revealed 3 bacteriophage groups that correlated with ICD-classified diseases diagnosed in the investigated patients: K29 *Gastritis and duodenitis*, K63 *Other diseases of intestine*, K29.6 *Other gastritis* (correlations defined by CI 0.95 and p<0.05) (Table 1).

ICD-correlated phage groups represented other (than species) taxonomic levels: *Clostridioides* prophages, *Tequatrovirus*, *Inovirus*. The first identified group of phages directly indicates a link to pathogenic bacteria that cause severe infections of the gastrointestinal tract (*Clostridioides difficile*), even though bacteria of that group were undetectable in the metagenomic sequencing of 16S RNA coding regions. This again suggests the role of phages as biomarkers with the potential to improve detection of bacterial causative agents of health disorders. Two latter phage groups are common phages that prey on *E. coli*, thus also suggesting either an increased fraction of aggressive pathogenic strains of that bacterial group or strongly disturbed conditions within the stomach (and possibly other parts of the gastrointestinal tract) that allow for overgrowth of normal bacterial biota.

Table 1. Bacteria and bacteriophage groups correlated with International Classification of Diseases (ICD)-classified diseases diagnosed in the investigated patients. The bacterial fraction of patients’ microbiome was identified by 16S rRNA sequencing of the mucosal bacterial fraction from stomach biopsies. Bacteriophage groups in the patients’ microbiomes were identified by shotgun sequencing of the mucosal viral fraction from stomach biopsies. ICD classification was carried out by qualified physicians following examination. The p-value was calculated using the z-test function from the statsmodels Python 3.6 library.

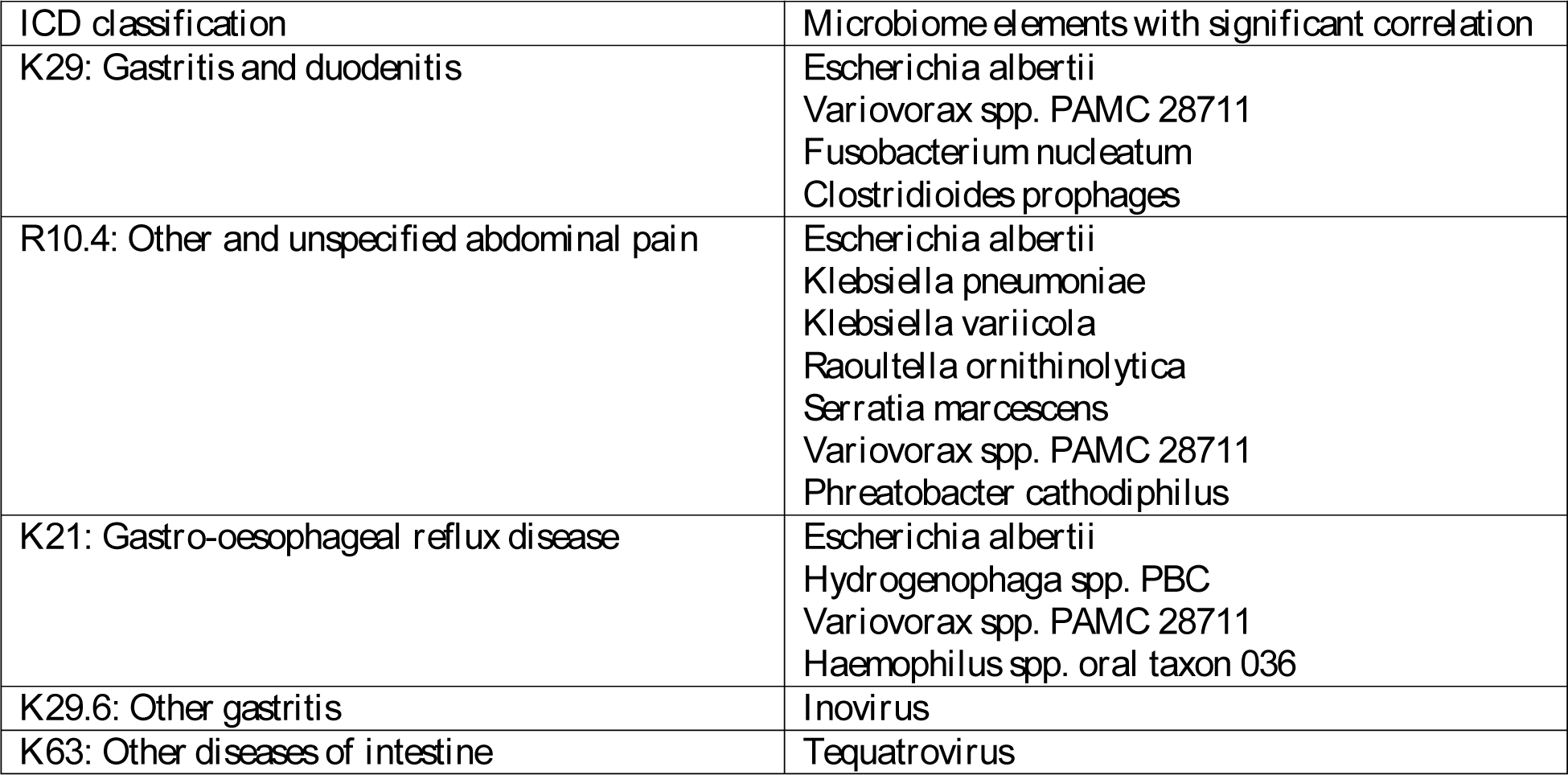

### Effects of human genotype on bacterial and phage microbiome composition (three-way SNP-phage-bacteria effects)

To identify tripartite correlations between human host genetic determinants, bacteriophages, and bacterial elements of the stomach microbiome, patients’ genotypes were analyzed by genotyping. The patients were genotyped with an AmpliSeq panel targeting 212 SNPs with confirmed links to human gastrointestinal, metabolic, or immunological disorders (a full list of the SNPs is presented in Table S4). Some SNPs were found to have a profound effect on bacterial and phage microbiome composition. A full list of SNPs that correlated with r>0.5 and p<0.05 with microbiome components is presented in Table S5. Among observed correlations, intercorrelated bacteriophages specific to *Staphylococcus* spp., also correlating with their hosts (Table S2), were significantly associated with two SNPs located in peptidoglycan recognition protein coding gene: rs3006438 (G/A) (PGLYRP4) and rs3014859 (A/G) (PGLYRP4), and with one SNP located in interleukin 22 coding gene rs2227473 (C/T) (IL22), even though *Staphylococcus* sp. was not found associated to these SNPs. Moreover, rs3006438 (PGLYRP4) presence was found to be significantly (p<0.05) correlated with microbiome diversity change in the patients’ microbiome; its presence of rs3006438 in the PGLYRP4 gene was linked to a 17.2% decrease in the Shannon index.

In the first part of this study, we observed that the presence of *Escherichia* strongly correlated (R>0.9) with the occurrence of the *Lederbergvirus* genus that includes coliphages (among other enterobacteria phages). A common SNP that correlated with enterobacteria phages *Lederbergvirus, Oslovirus,* and *Pankowvirus* and at the same time with *Escherichia* sp. and *Shigella* sp. was found to be the rs199883290 (duplicated C) allele in the *NOD2* gene. The gene is involved in cellular activity of the human immune system, as well as autophagy. IL1B allele, rs189235692 (A/G), was significantly correlated with *Escherichia*- and *Shigella*-specific phages like *Pankowvirus*, *Lederbergvirus*, and *Oslovirus*, and at the same time with their hosts *Escherichia* and *Shigella*. IL1B gene responsible for coding interleukin 1 beta, a crucial immunological component involved in the inflammatory response. Another human genetic variant that correlated with *Oslovirus*, *Pankowvirus*, *Lederbergvirus*, and *Escherichia sp.* and *Shigella sp.* was the rs115198942 (C/T) variant located within the intron region of IL23R. Moreover, the same correlations were found for another IL23R SNP (rs41313262). Interleukin 23 is also a pro-inflammatory cytokine; thus, its expression and regulation may affect immune responses to microbes.

In this study, the presence of specific bacteriophages was however predominantly found to correlate not with their respective bacterial hosts, but instead with the occurrence of SNPs in the human hosts of the studied microbiomes. For instance, *Inovirus*, primarily hosted by *Escherichia* spp., showed a correlation with the SNP rs2102654780 located in the IL23R region. Similarly, *Felsduovirus*, predominantly infecting *Klebsiella* and to some extent *Salmonella*, was associated with the SNP rs568408 (G/A) in the IL12A region, implicating potential effects of these pro-inflammatory immunological factors on bacteriophages. On the other hand, *Kuttervirus*, primarily associated with *Escherichia* and *Salmonella*, demonstrated a link to SNP rs4684677 in the *GHRL* gene. This gene encodes the preproghrelin protein, the precursor to ghrelin, a hormone chiefly produced by the stomach. This hormone regulates several physiological processes, including appetite and energy balance – hence being called the “hunger hormone”. This indicates that metabolic factors also shape the phageome composition. Interestingly, bacteriophages specific to *Escherichia* from the *Teseptimavirus* group, although not directly correlated with the presence of their bacterial host in this study, were instead found to correlate with SNPs in the *IL1B*, *TLR1*, and *TLR10* genes encoding potent immunological factors: interleukin 1 beta, and toll-like receptors 1 and 10. A similar trend was observed with *Lambdavirus*, *Tequatrovirus* and some staphylococcal bacteriophages, where correlated with SNPs in genes encoding immunological factors including IL6, NOD1, TLR10 and LTA. These findings suggest a significant influence of human genetic variability on the bacteriophage population, contributing to the complex interplay of genetic factors in shaping the microbiome.

In general, the most common variants that correlated with the occurrence of bacterial viruses were rs568408 (G/A) in the *IL12A* gene (13 bacteriophage groups) and rs3093664 in the TNF region (11 bacteriophage groups). When all individual SNPs in a single gene are considered, the gene polymorphisms more frequently associated with the presence of bacteriophages include IL23R (63 times), IL1B (45 times), IL22 (43 times), LTA (37), TLR10 (34 times) and IL6 (25 times), as calculated based on the number of identifications across all bacteriophage groups in all patients. All correlations between bacteriophage elements of the gastric microbiome, their hosts, and SNPs detected in the patients’ genome can be found in Supplementary Tables S2 and S5.

## Discussion

### 1. Triadic dynamics of gastric bacterial microbiome, phageome, and host genotype

Bacteriophages existing in the gastrointestinal tract interact with their hosts in many ways. Although the dynamics of bacterial viruses’ interactions with their hosts are not yet fully understood, certain patterns can be discerned (Lepage et al., 2013; Li et al., 2023; Shuwen & Kefeng, 2022). Beyond predator-prey dynamics, interactions between phages and bacteria suggest that lysogenic phage infection can have an impact on the bacterial host (Zuppi et al., 2021). All bacteriophage-bacteria interactions can be categorized into groups due to their complexity. Models in the first category are characterized by modest bacterial and phage population fluctuation and diversity. They include the Piggyback-the-Winner (PtW) model, which is characterized by mutualistic interactions that take place in the lysogenic life cycle, and the Arms-Race Dynamics (ARD) model, known by competition for survival driven by predator-prey interactions (Hampton et al., 2020; Knowles et al., 2016). The second set of models is distinguished by a significant and varied population of bacteriophages. These models are based on selection with negative frequency dependence. They include Fluctuating Selection Dynamics (FSD), which are unfavorable in an environment where numerous bacterial species fight for resources due to the fitness costs involved with the development of phage resistance mechanisms in bacteria, and the Kill-the-Winner (KtW) model, which happens when phage predation regulates the abundance of the “winning” bacterial species, i.e., the most competitive (Avrani et al., 2012; Thingstad, 2000). All these models demonstrate that bacteriophages and bacteria are strongly interdependent when existing in the human gastrointestinal tract. In this study, the stomach microbiome was investigated, with the focus on tripartite associations between the bacterial fraction, stomach bacteriophages, and the genetic background of a mammalian host. Our goal was to understand the associations among the components of the stomach microbiome, especially those linked to the ESKAPE and *Escherichia sp.* group of pathogens, renowned for their role in multi-drug resistant hospital infections. This goal included the human genetic component as a potential determinant of the microbiome composition.

Significant associations of bacterial groups were observed (Table S2), with strong correlations between *Escherichia* and *Shigella* (r=0.97), *Yersinia* (r=0.89), *Salmonella* (r=0.80), and *Enterobacter* (r=0.65). These can be partially explained by similar habitat preferences and physiological characteristics of these bacteria, which are members of the Enterobacteriaceae family. They share similar metabolic pathways and nutrient requirements, making them likely to thrive under similar conditions. Also, these microorganisms create specific niches by adapting the environmental conditions to their requirements. In this case, it is possible that their correlation has a mutualistic character; changes in the conditions initiated during the colonization by one microbial group influence other groups and create an opportunistic chance for growth for the others. The association between *Acinetobacter* species and *Helicobacter* can be explained similarly – through their influence on the nearby environment, they create a mutual opportunity to develop infection.

We further observed a strong correlation between *Klebsiella* and *Enterobacter* (r=0.95), *Yersinia* (r=0.86), *Providencia* (r=0.73), and *Morganella* (r=0.71). These associations might suggest an analogical resistance to the stomach’s harsh environment or co-evolution with a common human host immune response. Interestingly, *Acinetobacter* showed moderate correlations with *Helicobacter* (r=0.59) and *Enterococcus* (r=0.56). Although these genera are not a part of the same family and inhabit different ecological niches, their correlation might hint at indirect interactions or shared mechanisms of adaptation to the gastric environment. Our study revealed no significant correlations involving species of *Pseudomonas*, which could suggest its independent adaptation to the gastric environment or transient character of its participation in the microbiome. Additionally, we observed a moderate correlation between *Staphylococcus* and *Enterococcus* (r=0.55). Both genera are Gram-positive bacteria and are a part of the normal human microbiota, but they may include pathogenic species. Their correlation might indicate a shared ecological niche or similar mechanisms of survival in the stomach environment.

Within bacteriophage genera, the streptococcal phages *Moineauvirus* and *Brussowvirus* were significantly associated with each other, suggesting that presence of both groups was determined by the same bacterial hosts. Similarly, *Lambdavirus*, *Tequatrovirus*, and *Punavirus*, all targeting *Escherichia* species, showed a high correlation (r=0.78), possibly indicating co-feeding and population control of susceptible bacteria, and/or shared mechanisms of adaptation to *Escherichia*’s ecological niche, even though no association of these phage groups directly with their bacterial host was detected.

We expected significant associations between presence of bacteriophages and their specific hosts; however, this was not found as a universal principle. On the one hand, strong correlations between *Staphylococcus* and staphylococcal phages (*Biseptimavirus*, *Dubowvirus*, *Phietavirus*, and *Peeveelvirus*) were observed. Moreover, SNPs linked to the presence of both *Staphylococcus* and staphylococcal phages were identified; they were located in the *PGLYRP4* gene coding for peptidoglycan recognition protein 4, which plays a role in innate immunity and has been shown to have bactericidal activity against some Gram-positive bacteria (De Marzi et al., n.d.; Lu et al., 2006), and in the *IL22* gene that codes for the strongly pro-inflammatory cytokine interleukin 22. Also, a strong correlation between the presence of *Escherichia* and some relevant bacteriophages, particularly those belonging to *Pankowvirus*, *Lederbergvirus*, or *Ceceduovius*, was observed. Here also SNPs associated with both bacteriophages and their host were identified, including those located in the *NOD2* and *IL1B* genes, both coding for immunoreactive proteins, playing a role in pathogen recognition and stimulation of immune responses. Variations in the *NOD2* gene have been associated with several inflammatory disorders including Crohn’s disease and Blau syndrome. It is involved in the recognition of bacterial compounds and initiation of an immune response (Al Nabhani et al., 2017; Bauer et al., 2023; Ripoll et al., 2023; Walia & Mujahid, 2023). The rs115198942 (C/T) variant located within the intron region of the *IL23R* gene is correlated with both *Lederbergvirus* and *Escherichia* hosts. *IL23R* encodes the interleukin 23 receptor, a key player in immune responses and inflammation. The variant’s impact on IL-23 receptor expression and regulation could influence immune responses to microbes. Further research is needed to fully understand the functional implications of this variant (Álvarez-Salamero et al., 2020; Moschen et al., 2018; Sun et al., 2020). Of note, according to RegulomeDB, SNPs rs3014859 in *PGLYRP4* and rs199883290 in *NOD2* (detected in this study) have been scored 1f, thus demonstrating regulatory functions in the regions.

On the other hand, despite the confirming examples listed above, many more phage groups were not found to be clearly correlated with their bacterial hosts. Moreover, many phage groups were found to be correlated with specific genetic variants of human hosts (SNPs), particularly those linked to immunological functions and to important pro-inflammatory factors that are known to affect body responses to bacteria and viruses. These were for instance *Lambdavirus*, *Felsduovirus*, *Inovirus*, and *Teseptimavirus*. Bacteriophages from the *Kuttervirus* genus were associated with SNPs in a metabolic functions-related gene. A full list of the identified correlations is presented in Supplementary material (Tables S4 and S5). Since many phages within the stomach microbiome are associated with the human host genotype without parallel associations with bacterial hosts, it seems probable that the human body environment shapes phageomes by direct interactions. Thus, in addition to the expected effect on phage communities by shaping communities of their bacterial hosts, the human body seems to affect phages directly, selecting for phages that survive its specific selective pressure (Figure 3). Immunological factors were found to be the major tool of this pressure.

**Figure 3.**
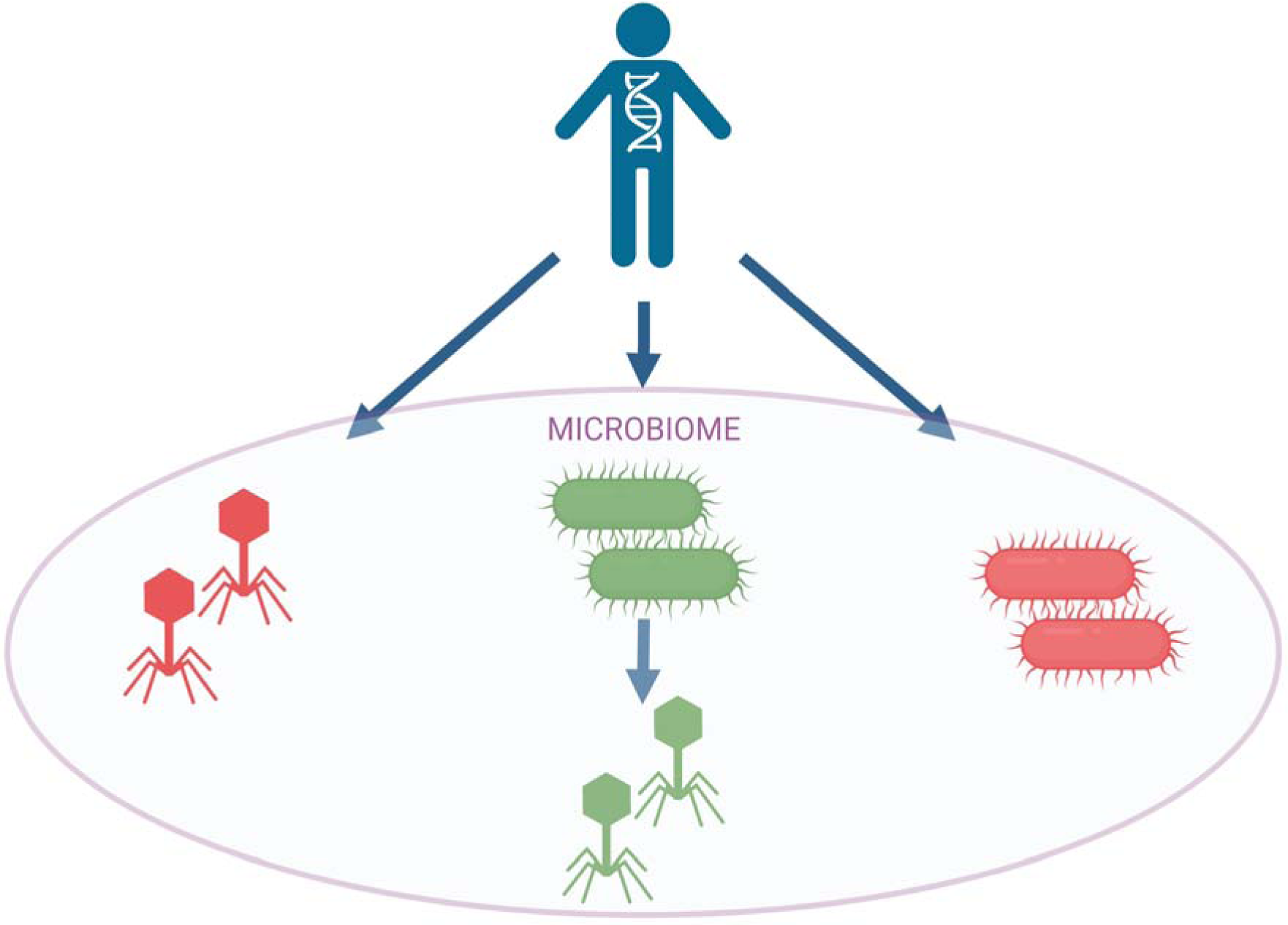
Human body shapes phage communities within body microbiome by shaping communities of their bacterial hosts (green) or directly selecting for phages that survive its specific selective pressure (red).

### 2. Implications for disease associations

Investigation of bacterial and phage components of the stomach microbiome in patients with ICD-classified disorders in the gastrointestinal tract (compared to control individuals) revealed both bacterial and phage elements of the microbiome that correlated with specific disorders. In bacteria, these correlations were expected since direct links between bacterial dysbiosis and multiple diseases have been reported (Carlson et al., 2018; McDonnell et al., 2021; Menni et al., 2020; Prehn-Kristensen et al., 2018). Associations between human diseases and bacteriophages have been previously described in the gut (not in the stomach). (Clooney et al., 2019; Jansen & Matthijnssens, 2023; Norman et al., 2015). Here we identified correlations of some gastrointestinal diseases and stomach phageome elements: *Clostridioides* prophages, tequatroviruses and inoviruses. The first group is related to a troublemaking pathogen, *Clostridioides difficile*. Two latter groups are common phages that parasitize *E. coli* but with very different life cycles. Inoviruses cause chronic infections of *E. coli* cells which lead to a continuous release of viral particles without host lysis, while tequatroviruses are obligatory lytic relatives of T4 phage. It remains unclear whether bacteriophages are causative agents or rather bioindicators of health disorders. The second possibility could include their host (*E. coli*), probably overrepresented in the bacterial microbiome of patients with gastrointestinal disorders. *E. coli* can be overrepresented as a result of an infection with pathogenic strains, or due to overgrowth of normally commensal strains in some disturbed conditions within the stomach (and possibly other parts of the gastrointestinal tract). Here the parallel correlation of the bacterial host was not detected. Alternatively, some phages (here, *Tequatrovirus* and *Inovirus*) can be promoted by specific conditions in the stomach, related to gastrointestinal disorders. These could be for instance increased pH or invalid production of digestive enzymes. Nevertheless, this issue calls for further investigation.

PCA showed two clusters that significantly differ when considering the overall microbiome composition. Pathogens such as *Acinetobacter* and *Helicobacter* had a strong influence on clustering (there was also a statistically significant correlation of occurrence between these pathogens, Table S2). However, viruses specific to *Lactococcus* and *E. coli* were also linked to different clustering of our samples. Population control by *Lactococcus*-specific viruses can affect the balance of the gastric microbiome. Possibly, deficiency of these bacteria through the over-presentation of specific bacteriophages may lead to the availability of receptors for potentially pathogenic microorganisms such as – also identified – *Acinetobacter* and *Helicobacter* specie. We hypothesize that the imbalance revealed by PCA shows how favorable conditions for the development of infection by some bacterial pathogens are created.

Some important limitations need to be considered for this study. First, bacteriophage identification is still restricted by the content of relevant databases. Phageomes, both environmental and body-related ones, still seem to be only partially recognized due to the extreme diversity of bacteriophages. Second, importantly, correlations do not indicate causation; rather, they provide a foundation for further inquiry into the investigated interactions. It means that in this study we cannot establish whether microbial groups that show significant correlations cause specific physiological effects or rather they are biomarkers. Mechanisms underlying identified correlations need to be investigated in further studies.

The most evident correlations observed here, however, indicate the immunological system as the major factor affecting phageomes in human bodies. Phage correlations with bacterial hosts were expected and observed, but a significant proportion of the correlations were bacterial host independent. Specifically, we identified some phage groups that correlated with specific human SNPs, without co-correlating susceptible bacteria. This is in line with the increasing body of evidence that phages in human bodies are strongly affected by the human immune system, which shapes phage activity and survival. Effects of the human organism on the phageome seem to be widely exerted by its immunity directly, not only by shaping the bacterial community serving as hosts for propagation of bacteriophages.

## Data accession

Sequencing data is available in Sequence Read Archive under the PRJNA934363 accession number.

## Acknowledgements

Project funded by the National Science Center grant UMO-2018/29/B/NZ6/01659.

